# Development and characterization of a high fat diet-streptozotocin induced type 2 diabetes model in nude athymic rats

**DOI:** 10.1101/2021.11.27.470203

**Authors:** Xuetao Sun, Sara S Nunes

## Abstract

People with diabetes mellitus (DM) are at an increased risk for myocardial infarction (MI) than age matched people without DM. However, assays for pre-clinical therapy are performed in animal models of ischemia that lack the co-morbid conditions present in patients with MI, such as DM. This contributes to the failure to translate pre-clinical trials results to the clinic. Thus, to increase the clinical relevance of xenograft studies in pre-clinical models, it is important to have a DM model in animals that are immunodeficient. Here, we developed a type 2 diabetes mellitus (T2D) model in nude athymic rats using high-fat diet and streptozotocin (HFD-STZ). Nude athymic rats were randomized into a control group (normal chow) or a HFD (45% fat, 20% protein and 35% carbohydrate)-STZ group. STZ (35 mg/kg i.v.) or vehicle was administered 8 weeks after HFD feeding started. Assessments were done longitudinally and at week 9 (endpoint). The HFD-STZ group showed mild hyperglycemia pre STZ administration (7.7 ± 0.3 mM *vs* 5.8 ± 0.2 mM in control) by week 8. In addition, plasma insulin levels were increased and the HOMA index was 2.5-times higher in the HFD-STZ. The HFD-STZ group showed increased fasting (147%) triglycerides. After STZ-administration, blood glucose levels increased substantially (23.6 ± 1.4 mM *vs* 5.5 ± 0.3 mM in control). The HFD-STZ treated animals also showed increased left ventricular wall thickness, cardiac hypertrophy and fibrosis, reduced cardiac function compared to normal chow control. In line with the HFD-STZ model in immunocompetent rats, the HFD-STZ treatment of athymic rats recapitulates key features of T2D, including aspects of established clinical diabetic cardiomyopathy and should be suitable for xenograft studies in the context of T2D.

## Introduction

Diabetes mellitus (DM) is a major metabolic disorder with increasing prevalence and has been considered a global epidemic. Cardiovascular disease (CVD) is the leading cause of morbidity and mortality in patients with diabetes, accounting for an estimated 80% of all diabetic deaths in North America^1^, including those from myocardial infarction (MI) and heart failure. The massive cardiomyocyte loss due to acute MI in the infarcted region is irreversible. Therefore, development of strategy to restore cardiac function of the diabetic MI heart may present a great therapeutic potential. Recent advances in cell transplantation suggest that cell replacement therapy may represent a promising approach to repair injured myocardium and improve cardiac function^2–5^. Animal models are widely used to study the underlying mechanisms of DM and to develop drugs for treating DM. There are two type of DM, type-1 (T1D) and type-2 (T2D). T2D is the predominant form, accounting for 90% of cases worldwide^6^. Many rat models have been established for research of T2D^7–9^. However, there are no reports of T2D models in immunocompromised rats, which are commonly/widely used for cell replacement therapy via transplantation of human cells^3,5,10,11^.

To increase clinical relevance in xenograft studies, it is important to have a T2D model in animals that are immunodeficient. In the present study, we adapted a well-established rat model of T2D induced by feeding a high fat diet and performing a single dose STZ injection^7^ in nude athymic rat and monitored the development of insulin resistance and hyperglycemia. Given our interest in cardiac regeneration, we also characterized the hart structurally and functionally for the hallmarks of diabetic cardiomyopathy. We describe that RNU nude rats were fed a HFD for 8 weeks developed the hallmarks of T2D: hyperinsulinemia, insulin resistance, and elevated Homeostatic Model Assessment for Insulin Resistance (HOMA-IR) score with a slight but significant increase in blood glucose levels. An additional increase in blood glucose levels was observed after a low dose STZ injection. Importantly, to ensure that diabetes-induced myocardial dysfunction was already established prior to MI induction and treatment (cell transplants), we confirmed the presence of the cardinal features of diabetes-induced myocardial disfunction, including: presence of cardiac hypertrophy at different scales (increased heart weight/tibial length, left ventricle (LV) wall thickness, and cardiomyocyte hypertrophy), increased fibrosis (assessed by picrosirius red staining) and an 18% decrease in vessel density compared to age matched, non-diabetic RNU nude rats at the 9-week time point. Echocardiographic assessments also revealed a significant increase in left ventricle internal diameter end systole (LVIDs) and a 5% decrease in fractional shortening. Our data indicates that the HFD-STZ treatment of athymic rats recapitulate key features of human T2D.

## Materials and methods

### Animal

Male athymic rnu/rnu rats (age 6-7 weeks, Charles River) weighing 160-200 g were used. All rats were housed in plastic cages in standard laboratory conditions (20°C, 56% RH). Rats were maintained under a 12-h light/dark cycle (6 am/ 6 pm) and had ad libitum access to food and water. All animal studies were approved by the Animal Care Committee of University Health Network (Toronto, Ontario, Canada) and conducted in accordance with the Guide for the Care and Use of Laboratory Animals published by the National Institutes of Health and Canadian Council on Animal Care guidelines.

### Induction of T2D by HFD-feeding and streptozotocin at low dose

The normal chow consisted of 3.1 Kcal/g, comprising 17% calories from fat, 25% from protein and 58% from carbohydrate (Teklad, Envigo, Madison, WI, USA). The HFD consisted of 4.7 Kcal/g, comprising 45% calories from fat, 20% from protein and 35% from carbohydrate (Research Diets, New Brunswick, NJ, USA). After 8-week HFD feeding, rats were injected intravenously via tail vein with a single low dose of streptozotocin (STZ) (35 mg/kg) (Sigma). One week after STZ injection, rats with diabetes showing hyperglycemia (blood glucose ≥15 mmol/L (270 mg/dl)) were considered diabetic.

### Assessment of insulin resistance

Homeostasis model assessment-insulin resistance (HOMA-IR) was used to assess β-cell function and insulin resistance (IR) from basal glucose and insulin. The fasting blood glucose concentration from the tail vein was measured using a OneTouch blood glucose monitoring system (LifeScan, Malvern, PA, USA). The plasma insulin concentration was determined using rat insulin enzyme-linked immunosorbent assay kit (Elisa) (Cat.10-1250-01, Mercodia, Uppsala, Sweden). The results were expressed as μU/mL of plasma insulin. HOMA-IR was calculated as fasting glucose (mmol/L) × fasting insulin (μU/mL)/22.5 as described previously^12,13^.

### Analysis of insulin sensitivity

Insulin tolerance test was performed to assess the insulin sensitivity. After fasting for 6h, rats were injected intraperitoneally with 1 U/kg human insulin (Sigma). Blood glucose from the tail vein was measured using a glucometer at the desired time (0, 15, 30, 60, 90 and 120 min) after the insulin administration.

### Cardiac Blood Panel

A cardiac blood panel was run at week 8 to measure levels of high-density lipoprotein (HDL)-cholesterol, low-density lipoprotein (LDL)-cholesterol, total cholesterol and triglyceride. Animals were fasted for 6 h and blood was taken from the saphenous vein and collected in EDTA-coated tubes. Tubes were centrifuged for 5 min at 500 rcf to separate blood plasma. Blood plasma was analyzed with a Beckman Coulter analyzer (The Centre for Phenogenomics).

### Cardiac function

Echocardiography was performed at day 0 (baseline), before STZ injection (week8) and after STZ injection (week 9). The left-ventricular end diastolic dimension (LVEDD) and the left-ventricular end systolic dimension (LVESD) were measured using a GE Vivid 7 Dimension with a 10S (10 MHz) pediatric probe, and fractional shortening (FS) was calculated by the equation: FS = 100 x (LVEDD – LVESD)/LVEDD^5,14^. LVEDD and LVESD are expressed in mm.

### Histology and immunocytochemistry

Body weight, heart weight, tibial length were measured^15^. Dissected hearts were fixed in 10% neutral-buffered formalin (Sigma). Then the hearts were cut into 2-mm thick transverse slices using a matrix slicer (Zivic Instruments, Pittsburgh, PA) before standard paraffin-embedding. Five-micrometer sections were then cut and stained with hematoxylin-eosin (Sigma) for quantification of left ventricular wall thickness or Picrosirius red for quantification of fibrosis^5^.

Immunostaining was performed with antibodies monoclonal mouse anti-troponin T (1:100, Fisher Scientific) and Alexa488 donkey anti mouse (Thermofisher) with nuclei counterstained with Hoechst 33342 (1:1000, Sigma). To detect CD31/PECAM (1:200, Novus), biotinylated goat anti rabbit secondary antibody (Vector Labs) was used in conjunction with the Vectastain Avidin/Biotin Complex (ABC) Kit (Vector labs) followed by alkaline phosphatase (Vector Red) or HRP/DAB (Vector Labs, Burlingame, CA, USA). The sections were counterstained with Haematoxylin (Sigma) to detect nuclei. Sections were mounted using vectashield antifade mounting medium (Vector labs) or Fisher chemical permount mounting medium (Fisher Scientific). Fluorescence images were captured using Nikon DS-Ri1-U2 USB camera attached to Nikon Ti Eclipse microscope using Nikon NIS elements BR (3.22.11). Confocal fluorescence images were captured using Olympus Fluoview 1000 Laser Scanning Confocal Microscope. Images of histological microscope slides were scanned by a Leica AT2 Scanscope and analysed using ImageJ (1.49v). Cell surface area was measured by rhodamine conjugated wheat germ agglutinin (WGA) (1:1500, Biotium) staining using ImageJ^5^. All quantification of histologic parameters was done in a blinded manner.

### Statistics

All animals were individually coded and maintained until all data were acquired and analyzed. Values are expressed as mean ± SEM unless otherwise stated. *P* < 0.05 was considered statistically significant. Data was analyzed using a two-way analysis of variance (ANOVA), where time and diet were independent variables, followed by a Bonferroni’s *post-hoc* test for multiple comparisons (SigmaPlot). A Student’s *t*-test was performed where diet was the only factor.

## Results

### Diabetes induction

Nude athymic rats were fed a HFD for 8 weeks when a low-dose injection of streptozotocin (STZ) (35 mg/kg, i.v.) was performed (**Fig. 1A**). Rats fed a normal chow served as healthy controls. HFD feeding for 8 weeks resulted in an insignificant increase in body weight than the normal chow fed control rats (**Fig. 1B**, week 8, *P* = 0.074). However, there was a modest weight loss after STZ administration (**Fig. 1B**, week 9).

**Fig. 1.**
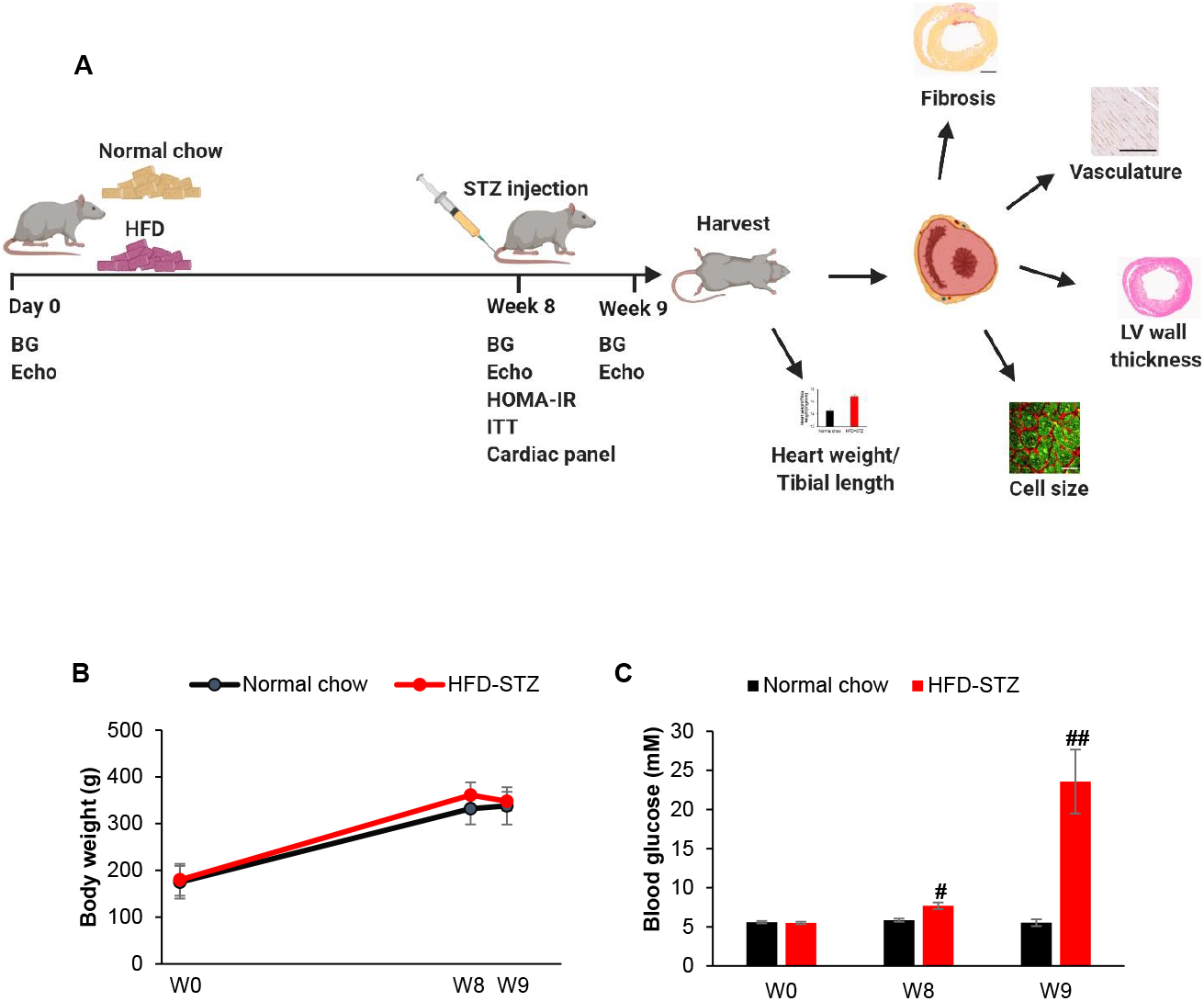
Time course of HFD-STZ induced hyperglycemia in nude athymic rats. (A) Schematic of the experimental protocol. BG, blood glucose. Echo, echocardiography. HFD, high fat diet. HOMA-IR, Homeostasis model assessment-insulin resistance. ITT, insulin tolerance test. LV, left ventricle. STZ, streptozotocin. (B) Body weight and (C) blood glucose were determined in normal chow control (n=7) and HFD-STZ animals (n=8). #, *P* < 0.05, ##, *P* < 0.001 *vs* normal chow.

At week 8, the blood glucose level was significantly higher in HFD-fed animals (7.7 ± 0.3 mM) compared to normal chow-fed controls (5.8 ± 0.2 mM) (**Fig. 1C**, *P* = 0.048). After STZ injection, the hyperglycemia in the HFD-STZ group increased significantly (**Fig. 1C**, week 9, 23.6 ± 1.4 mM), similarly to levels previously described in Wistar or Sprague-Dawley rats^7,9^.

To test if HFD feeding induced insulin resistance, we performed an insulin tolerance test at the 8-week time point prior to STZ injection. Fasting plasma insulin levels were significantly higher in HFD-STZ rats compared to normal chow-fed control (**Fig. 2A**). The degree of insulin resistance (IR) in the HFD-STZ group was significantly higher than in control, as shown by the assessment of HOMA-IR (**Fig. 2B**). Additionally, insulin sensitivity was reduced in HFD-STZ animals compared to control rats, as shown by the blood glucose curves (**Fig. 2C, D**).

**Fig. 2.**
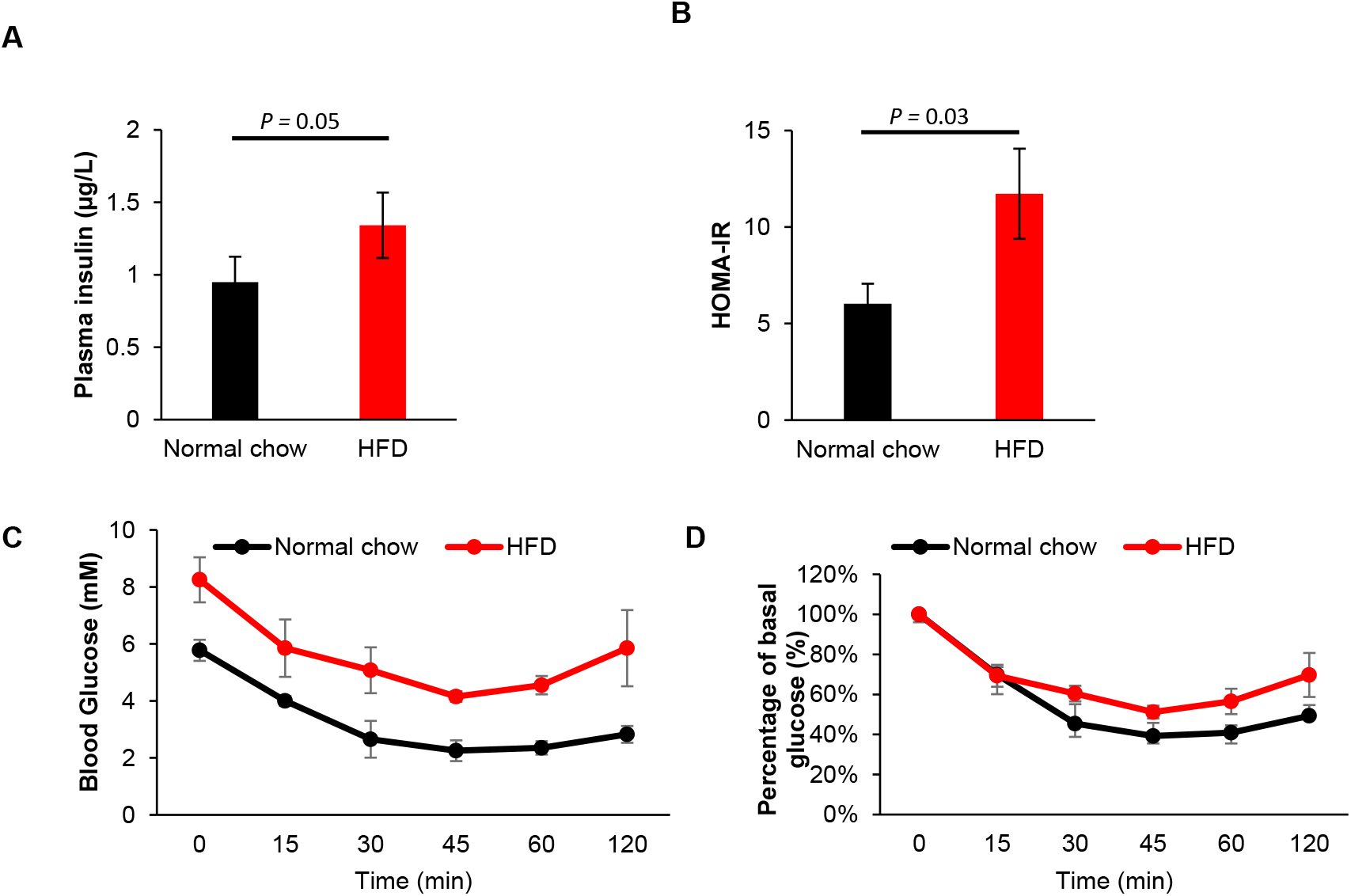
Insulin resistance and sensitivity analysis for athymic nude rats on normal chow or HFD at week 8. (A) The fasting plasma insulin levels. (B) HOMA-IR. HFD (n=8) or normal chow (n=7). (C) Blood glucose level during ITT. (D) Percentage of initial glucose level during ITT. *P*=0.011, HFD (n=4) vs Normal chow (n=4),

Assessment of blood lipids at week 8 revealed significantly increased HDL-cholesterol, total cholesterol and triglycerides in the HFD cohort relative to control (**Fig. 3**). No differences were observed for LDL-cholesterol between groups (**Fig. 3**).

**Fig. 3.**
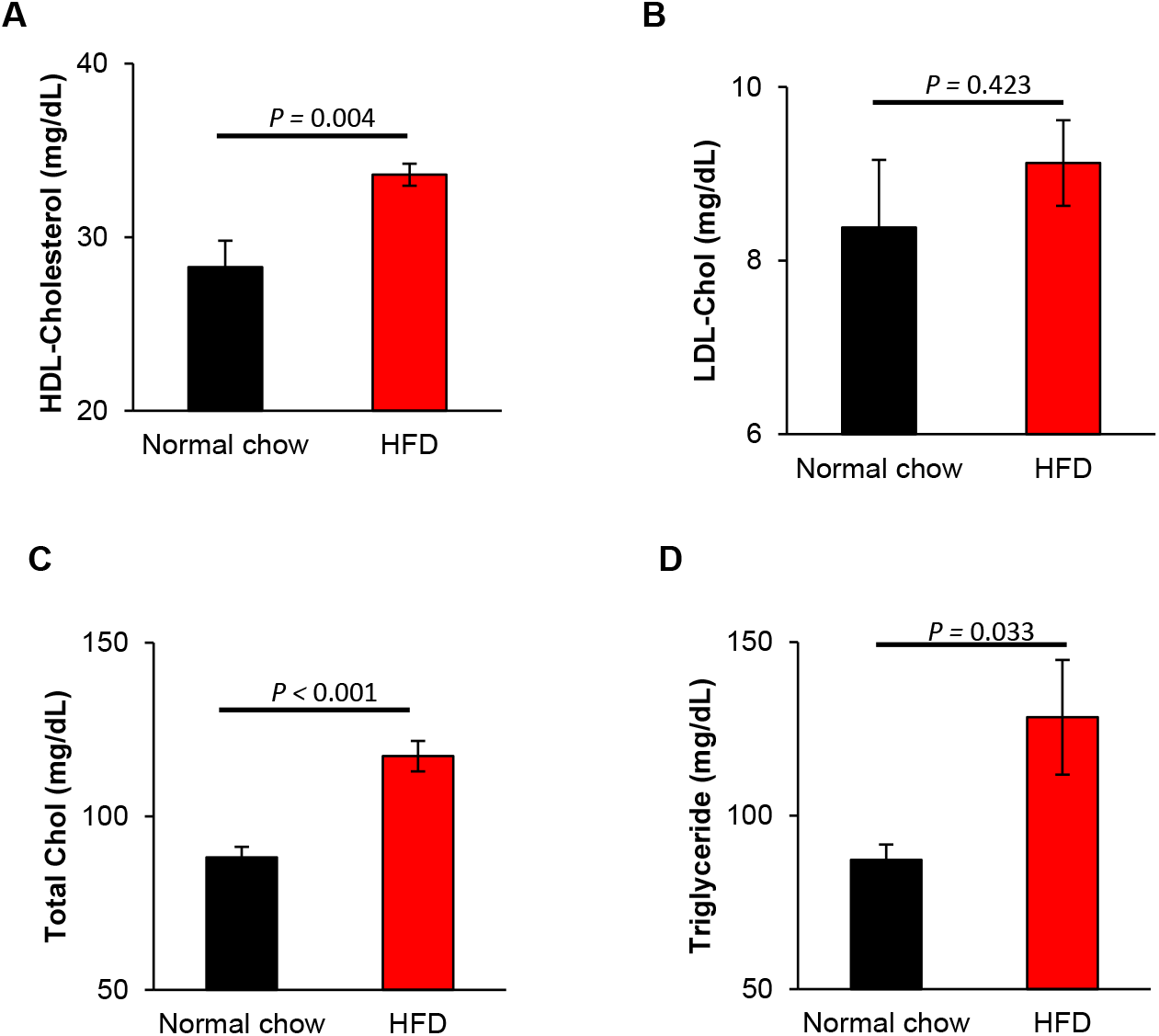
Cardiac blood panel for rats on HFD or normal chow at week 8. Rat blood samples were measured for (A) high-density lipoprotein (HDL)-cholesterol, (B) low-density lipoprotein (LDL)-cholesterol, (C) total cholesterol and (D) triglyceride.

### Analysis of cardiac function

Echocardiography was performed to assess changes in cardiac function longitudinally (**Fig. 4**). We observed that fractional shortening was significantly lower in HFD-STZ rats (43.7±2.2%) compared to healthy controls (48.7±1.0%) at week 9 (*P* <0.001, **Fig. 4A,**). In addition, at week 8 (prior to STZ injection), fractional shortening was already significantly lower in HFD fed animals (42.9±1.6%) compared to control (48.4±1.3%) (*P*<0.001, **Fig. 4A**). Despite an initial significant increase in left ventricular internal dimension in diastole (LVIDd) and left ventricular internal dimension in systole (LVIDs) in HFD rats at week 8, prior to STZ injection (**Fig. 4B, C,** week 8), after treatment with STZ there were no differences in LVIDd (8.2±0.3 mm) compared to control (7.9±0.5 mm) (*P*=0.051, **Fig. 4B**). LVIDs was still significantly higher in HFD-STZ rats (4.6±0.3 mm) compared to normal chow control (4.0±0.3 mm) (*P*<0.001, **Fig.4C**). These indicate that dietary manipulation alone could impact cardiac function in nude athymic rats.

**Fig. 4.**
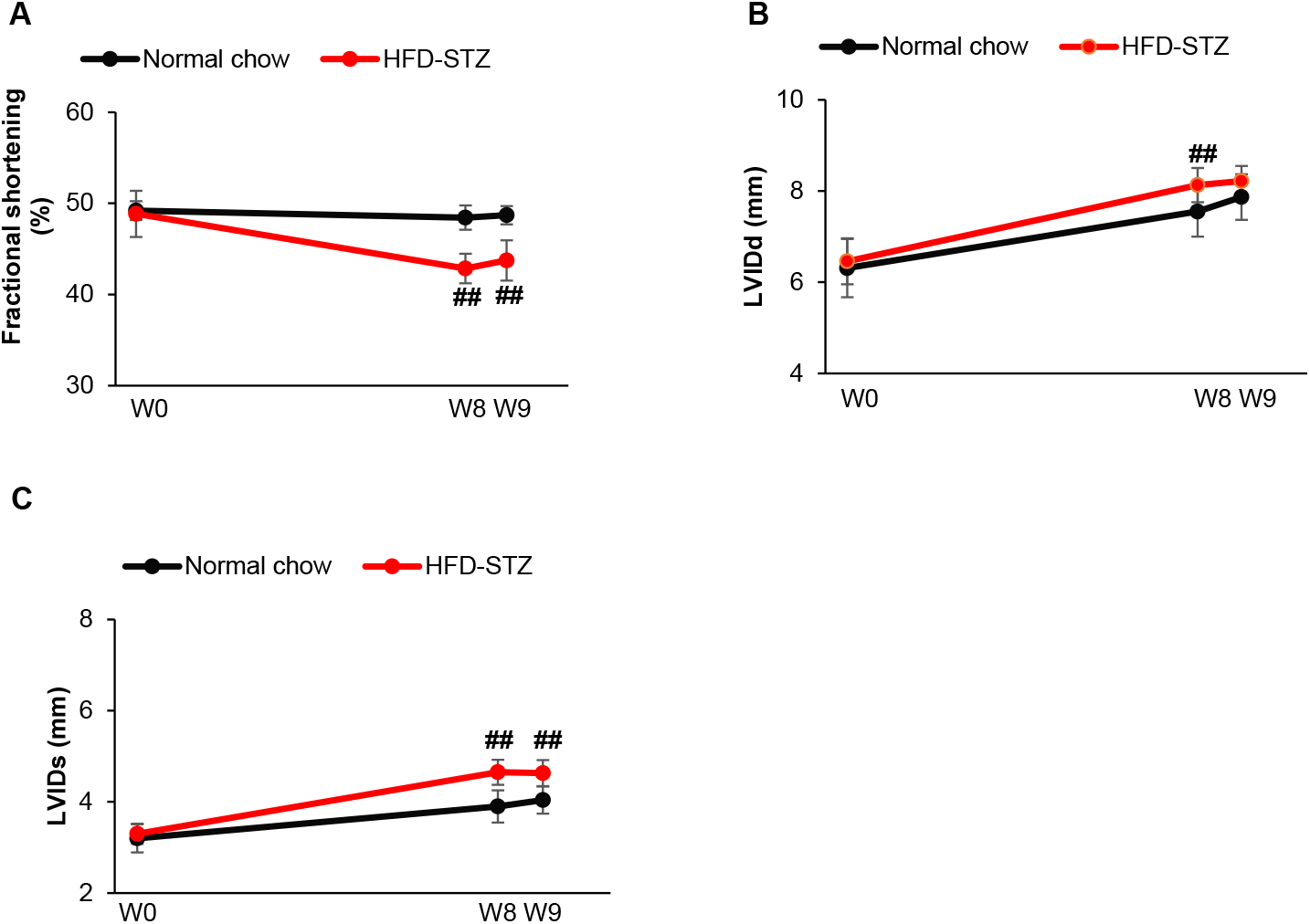
Cardiac function analysis by echocardiograph for rats on HFD or normal chow. HFD, n=7, normal chow, n=8. ##, *P* <0.001, *vs* normal chow.

### Morphological assessments of the heart

Assessment of the ratio of heart weight by tibial length indicated that HFD-STZ rats displayed significantly higher ratios compared to control rats (*P*=0.005, **Fig. 5A**), suggesting cardiac hypertrophy in HFD-STZ animals. Histological analysis confirmed the presence of hypertrophy, as shown by the increased left ventricular wall thickness in HFD-STZ animals compared to controls (*P*=0.005, **Fig.5B, C**). Assessment of the cardiomyocyte size via the measurement of surface area of WGA and cTNT stained sections^5^ showed that cardiomyocytes in the HFD-STZ rats were significantly larger compared to cardiomyocytes from control rats (*P*=0.001, **Fig. 5D, E**). We also examined whether fibrosis and/or microvascular rarefaction was present as these have been associated with DM^16,17^. Picrosirius red staining of cardiac sections revealed a significantly increased collagen deposition in HFD-STZ rats compared to normal chow control at week 9 (*P*=0.05, **Fig. 6 A, B**), indicating fibrosis. Myocardial vessel density was slightly reduced (18%) in the HFD-STZ rats (1743±67/mm^2^) compared to controls (2062±180/mm^2^), but not at a significant level (*P*=0.149, **Fig. 7**).

**Fig. 5.**
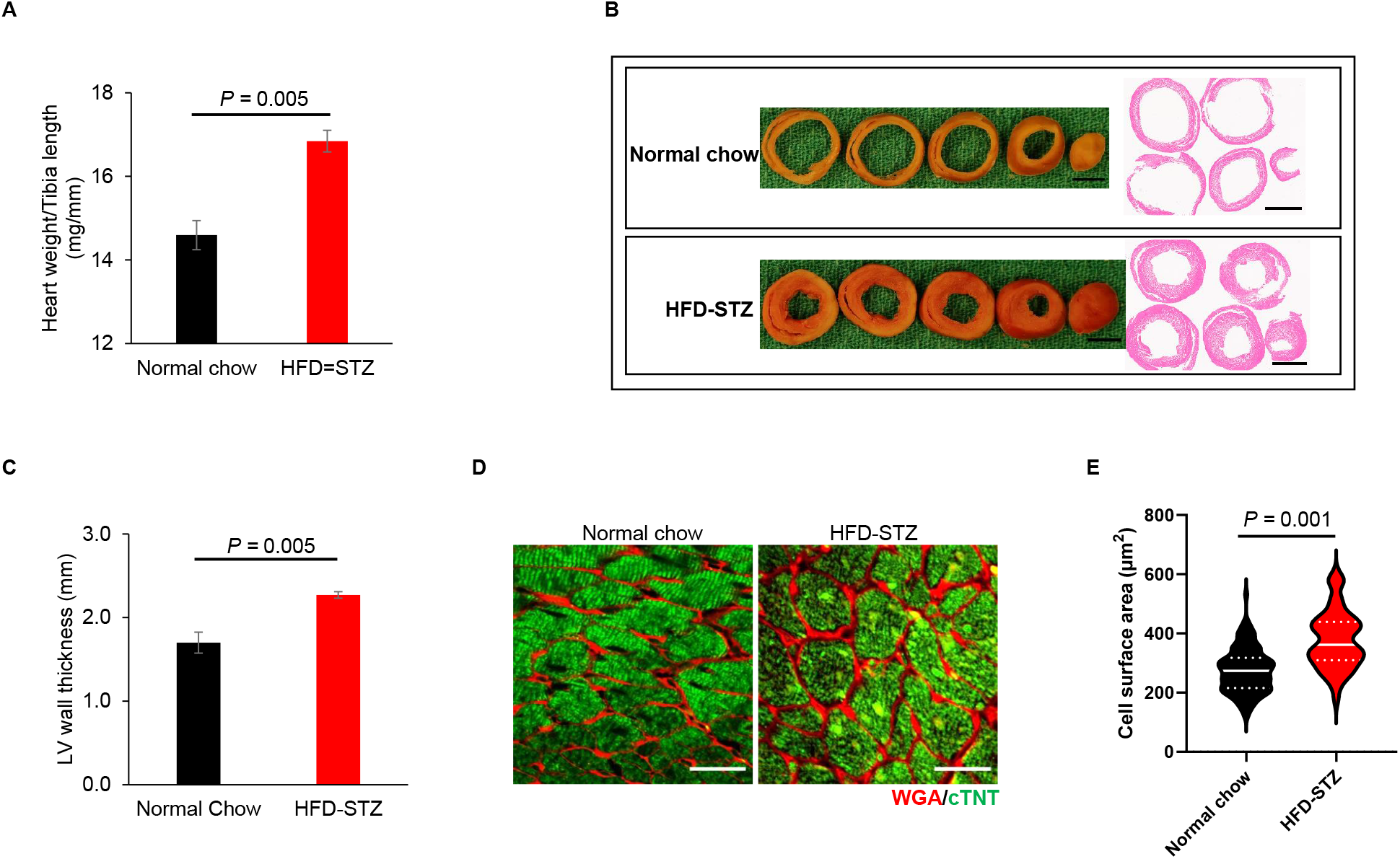
Cardiac structure of nude athymic rats on HFD-STZ or normal chow. (A) Ratios of heart weight/ and heart weight/tibia length. (B) Harvested hearts were sliced and embedded in paraffin for H&E staining. Scale bar, 500 mm. (C) Measurement of LV wall thickness. (D) Cardiomyocytes were stained with Alexa 488 conjugated cTNT and rhodamine conjugated wheat germ agglutinin (WGA). Scale bar, 50 μm. (E) Quantification of cardiomyocyte cell surface area.

**Fig. 6.**
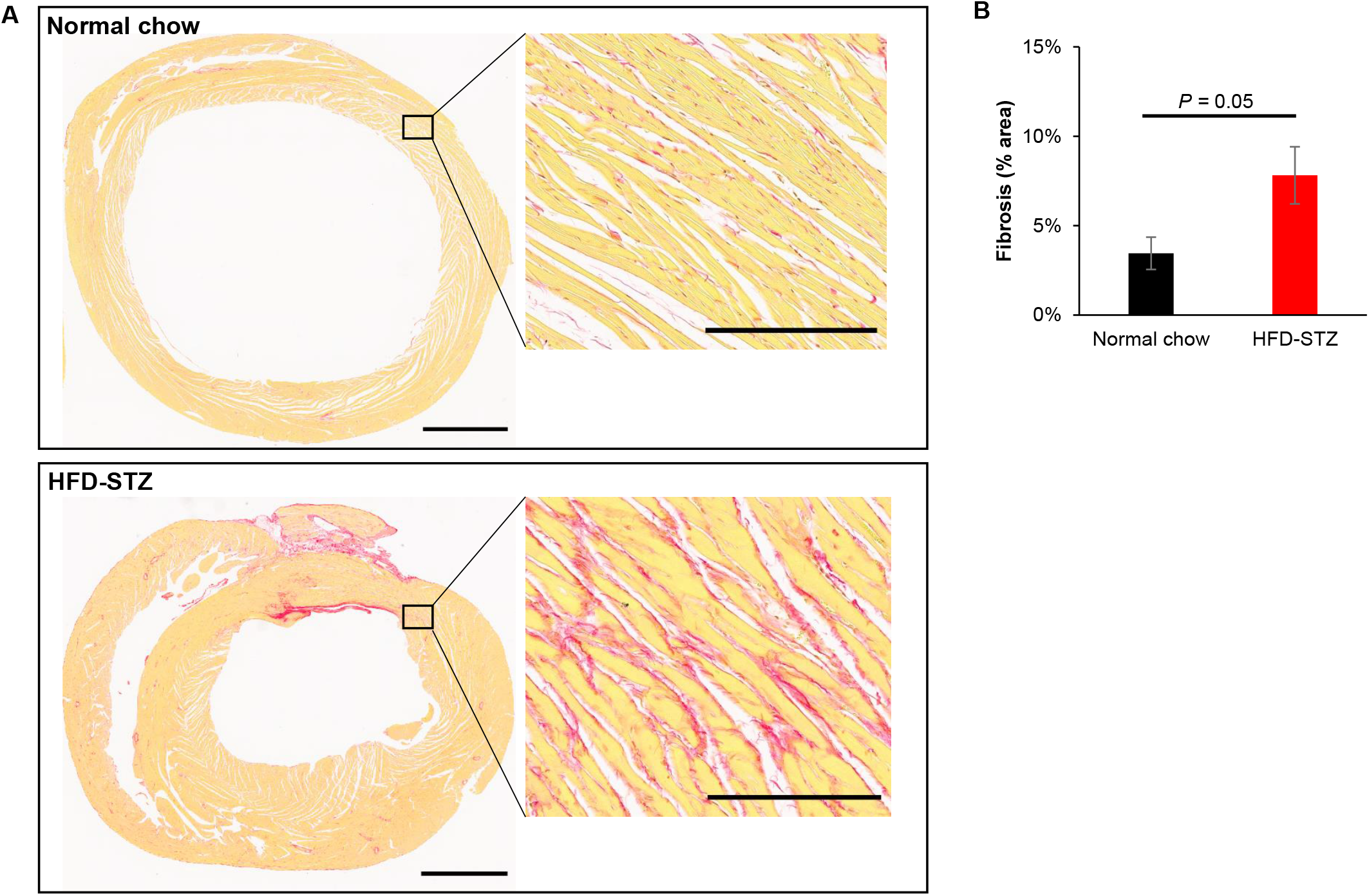
Cardiac fibrosis in nude athymic rats on HFD-STZ or normal chow. (A) Paraffin sections of heart at week 9 stained with picrosirius red. Scale bar, 2 mm, inset images,100 μm. (B) Quantification of fibrotic area. HFD, n=3, normal chow n=4, Student *t*-test.

**Fig. 7.**
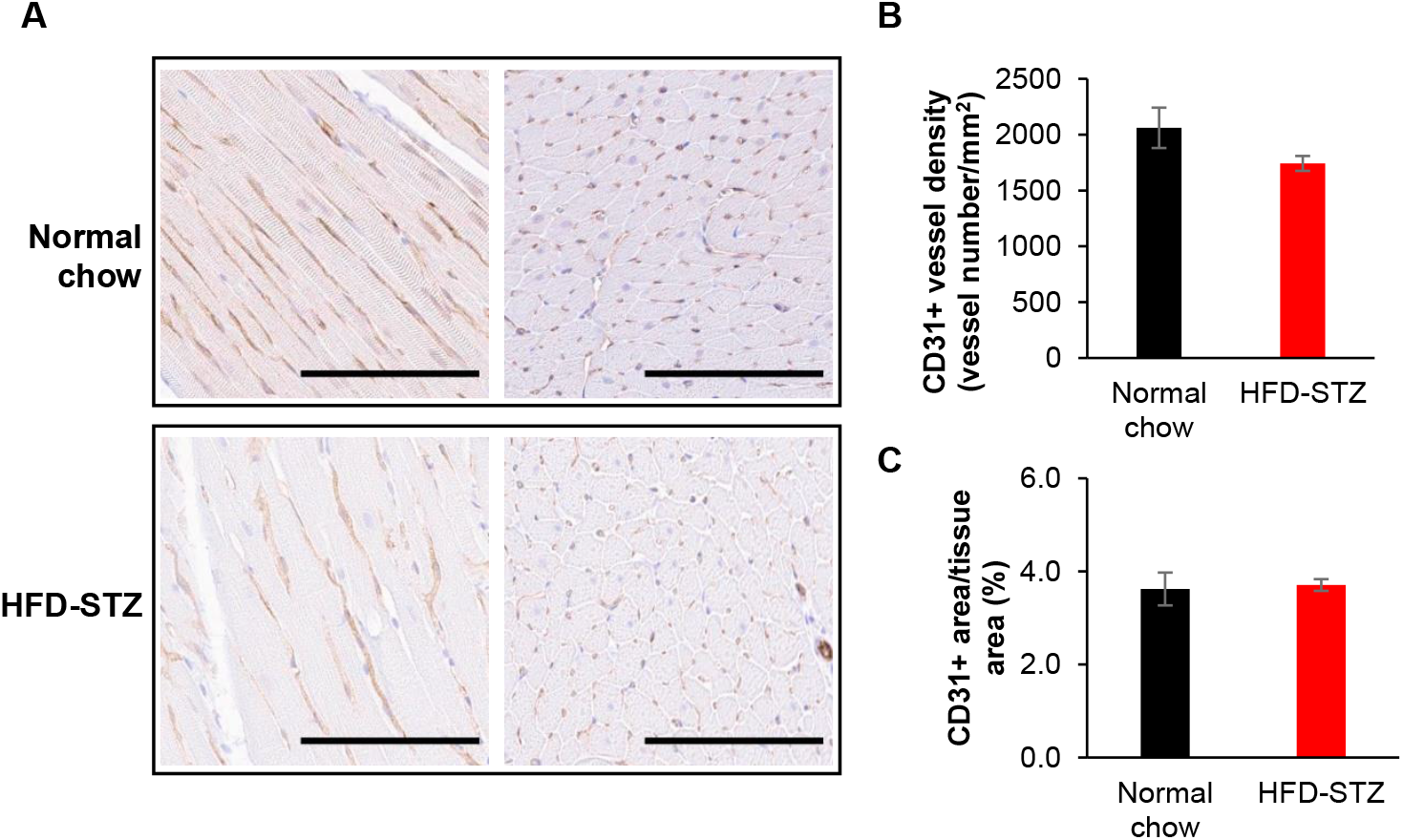
Cardiac vasculature in nude athymic rats on HFD-STZ or normal chow. (A) Paraffin sections of heart stained with CD31 with nuclei counterstained with hematoxylin. Scale bar, 50 μm. (B&C) Quantification of CD31+ vessel density (B) and (C) area. HFD, n=3, normal chow n=4.

## Discussion

In this study, we developed and characterized a T2D model using RNU nude rats that allow xenotransplantation studies. We showed that RNU nude rats display several hallmarks of T2D, including cardiac dysfunction. Animal models of T2D are valuable for studying the pathobiology of the disease. One well established and widely used T2D animal model is achieved by high fat diet feeding followed by STZ injection of rats. Though there are several versions of the HFD-STZ rat model, varying in length of HFD feeding and in STZ dose and number of injections, the protocols are similar: HFD feeding for short (2-4 weeks) or long (over 3 months) periods followed by STZ injection^7–9,18^. Given practical and financial considerations, our chosen regimen (8 weeks HFD feeding) was intended to be short while successfully showing key hallmarks of the disease, such as increased blood glucose levels, and insulin resistance.

There are great variations in the STZ treatments, which results in varied depletion levels of the insulin-producing β-cells. These variations include the dose (amount, frequency, time interval and the route of injection) and the fed/fasted state at time of injection^7–9,19^. Though there is no consensus on the STZ treatment in modeling of T2D using a HFD-STZ in rats, we elected to administer a low dose of STZ to decrease the potential of STZ toxicity^20^. As such, a single STZ injection (35 mg/kg) after 8 weeks of HFD-feeding was effective in inducing hyperglycemia. It should be noted that in the present study, STZ treatment led the RNU rats transit from an insulin-resistant state showing mildly elevated blood glucose levels to blood glucose levels of more established T2D within 1 week, which might not precisely mimic the chronological progression of the disease in humans and is therefore a limitation of this model.

It is well known that diabetes affects the heart and can lead to diabetic cardiomyopathy. The cardinal features of diabetic cardiomyopathy include myocardial fibrosis and hypertrophy with an initial cardiac diastolic dysfunction which subsequently progresses into systolic dysfunction and eventually into clinical heart failure^17^. Histologically, our data revealed that HFD-STZ induced myocardial thickening, cardiomyocyte hypertrophy and increased collagen deposition (i.e. fibrosis) compared to healthy controls by week 9. Assessment of myocardial microvascular density revealed absence of vascular rarefaction diabetic rats, which was unexpected. However, it is likely these features would take longer to appear as seen in chronic T2D. Overall, our findings are in line with the hallmarks of established clinical diabetic cardiomyopathy.

We also monitored several biomarkers widely used in the clinic for predicting heart disease risks including LDL cholesterol, total cholesterol, and triglycerides. Our data showed the elevated levels of total cholesterol and triglycerides. These changes are concurrent with the structural and functional cardiac changes. However, HDL-cholesterol, commonly known as the “good” cholesterol and associated with a lower risk of heart disease, was also significantly elevated. Similar result has been reported in diabetic rat^21^.

The immune system is a key element in T2D and contributes to its progression from prediabetes to fully established T2D in different ways^22^. Although RNU nude rats are immunocompromised, it is not surprising that they still showed several hallmarks of T2D as nude rats, although lacking T cells, have the full range of immune cells, including B and NK cells, monocytes, macrophages^23,24^.

However, there were several limitations on this study. First, sex is a component of rigour and reproducibility in animal study and female animals should be included in future to provide a more efficient and effective experimental design. Second, the age of the rats is also an important factor in modeling the HFD-STZ diabetes. We used 6-7 weeks as the initial age of the rats, similar to the majority of HFD-STZ rats reported using young rats^7,19,25^. Though the prevalence of T2D in children and youth is increasing, the disease is still much more prevalent in older age groups^26^. Meanwhile, β cell regeneration capacity may decline with age in mammals^27^. Therefore age should be also taken into consideration in future studies. Third, recovery of insulin production may need to be verified at longer time points, particularly if this model is to be used in studies involving transplantation of insulin-secreting beta-cells. Lastly, more analysis is required to identify the stage (early *vs* late) of T2D mimicked in this study.

In summary, we generated a HFD-STZ-induced T2D model using RNU nude rats that are amenable for xenograph studies. The characterization of the structural and functional cardiac changes associated with the progression of diabetes will allow us to study the compounded effects of T2D-myocardial dysfunction and ischemia (MI) in myocardial regeneration.

## Acknowledgements

This work was supported by a grant from the Canadian Institutes of Health Research (PJT153160) to S.S.N.

## Author contributions

X.S. was the main experimentalist. X.S. designed the experiments, analyzed the data and contributed to the writing of the manuscript. S.S.N. designed the experiments, coordinated the project, and contributed to the writing of the manuscript.

## References

1 Glass, C. E., Singal, P. K. & Singla, D. K. Stem cells in the diabetic infarcted heart. Heart Fail Rev 15, 581–588, doi:10.1007/s10741-010-9172-8 (2010).

2 Shiba, Y. et al. Human ES-cell-derived cardiomyocytes electrically couple and suppress arrhythmias in injured hearts. Nature 489, 322–325, doi:10.1038/nature11317 (2012).

3 Laflamme, M. A. et al. Cardiomyocytes derived from human embryonic stem cells in pro-survival factors enhance function of infarcted rat hearts. Nature biotechnology 25, 1015–1024, doi:10.1038/nbt1327 (2007).

4 Chong, J. J. et al. Human embryonic-stem-cell-derived cardiomyocytes regenerate non-human primate hearts. Nature 510, 273–277, doi:10.1038/nature13233 (2014).

5 Sun, X. et al. Transplanted microvessels improve pluripotent stem cell-derived cardiomyocyte engraftment and cardiac function after infarction in rats. Science translational medicine 12, doi:10.1126/scitranslmed.aax2992 (2020).

6 Chatterjee, S., Khunti, K. & Davies, M. J. Type 2 diabetes. Lancet 389, 2239–2251, doi:10.1016/S0140-6736(17)30058-2 (2017).

7 Reed, M. J. et al. A new rat model of type 2 diabetes: the fat-fed, streptozotocin-treated rat. Metabolism 49, 1390–1394, doi:10.1053/meta.2000.17721 (2000).

8 Srinivasan, K., Viswanad, B., Asrat, L., Kaul, C. L. & Ramarao, P. Combination of high-fat diet-fed and low-dose streptozotocin-treated rat: a model for type 2 diabetes and pharmacological screening. Pharmacol Res 52, 313–320, doi:10.1016/j.phrs.2005.05.004 (2005).

9 Zhang, M., Lv, X. Y., Li, J., Xu, Z. G. & Chen, L. The characterization of high-fat diet and multiple low-dose streptozotocin induced type 2 diabetes rat model. Exp Diabetes Res 2008, 704045, doi:10.1155/2008/704045 (2008).

10 Laflamme, M. A. et al. Formation of human myocardium in the rat heart from human embryonic stem cells. The American journal of pathology 167, 663–671, doi:10.1016/S0002-9440(10)62041-X (2005).

11 Fernandes, S. et al. Human embryonic stem cell-derived cardiomyocytes engraft but do not alter cardiac remodeling after chronic infarction in rats. Journal of molecular and cellular cardiology 49, 941–949, doi:10.1016/j.yjmcc.2010.09.008 (2010).

12 Matthews, D. R. et al. Homeostasis model assessment: insulin resistance and beta-cell function from fasting plasma glucose and insulin concentrations in man. Diabetologia 28, 412–419, doi:10.1007/BF00280883 (1985).

13 Mather, K. Surrogate measures of insulin resistance: of rats, mice, and men. Am J Physiol Endocrinol Metab 296, E398–399, doi:10.1152/ajpendo.90889.2008 (2009).

14 Mihic, A. et al. A Conductive Polymer Hydrogel Supports Cell Electrical Signaling and Improves Cardiac Function After Implantation into Myocardial Infarct. Circulation 132, 772–784, doi:10.1161/CIRCULATIONAHA.114.014937 (2015).

15 Yin, F. C., Spurgeon, H. A., Rakusan, K., Weisfeldt, M. L. & Lakatta, E. G. Use of tibial length to quantify cardiac hypertrophy: application in the aging rat. Am J Physiol 243, H941–947, doi:10.1152/ajpheart.1982.243.6.H941 (1982).

16 Hinkel, R. et al. Diabetes Mellitus-Induced Microvascular Destabilization in the Myocardium. Journal of the American College of Cardiology 69, 131–143, doi:10.1016/j.jacc.2016.10.058 (2017).

17 Jia, G., Hill, M. A. & Sowers, J. R. Diabetic Cardiomyopathy: An Update of Mechanisms Contributing to This Clinical Entity. Circulation research 122, 624–638, doi:10.1161/CIRCRESAHA.117.311586 (2018).

18 Hu, S. H., Jiang, T., Yang, S. S. & Yang, Y. Pioglitazone ameliorates intracerebral insulin resistance and tau-protein hyperphosphorylation in rats with type 2 diabetes. Exp Clin Endocrinol Diabetes 121, 220–224, doi:10.1055/s-0032-1333277 (2013).

19 Si, Y. et al. Infusion of mesenchymal stem cells ameliorates hyperglycemia in type 2 diabetic rats: identification of a novel role in improving insulin sensitivity. Diabetes 61, 1616–1625, doi:10.2337/db11-1141 (2012).

20 Szkudelski, T. The mechanism of alloxan and streptozotocin action in B cells of the rat pancreas. Physiol Res 50, 537–546 (2001).

21 Loai, S., Zhou, Y. Q., Vollett, K. D. W. & Cheng, H. M. Skeletal Muscle Microvascular Dysfunction Manifests Early in Diabetic Cardiomyopathy. Front Cardiovasc Med 8, 715400, doi:10.3389/fcvm.2021.715400 (2021).

22 Zhou, T. et al. Role of Adaptive and Innate Immunity in Type 2 Diabetes Mellitus. J Diabetes Res 2018, 7457269, doi:10.1155/2018/7457269 (2018).

23 Festing, M. F., May, D., Connors, T. A., Lovell, D. & Sparrow, S. An athymic nude mutation in the rat. Nature 274, 365–366, doi:10.1038/274365a0 (1978).

24 Rolstad, B. The athymic nude rat: an animal experimental model to reveal novel aspects of innate immune responses? Immunol Rev 184, 136–144, doi:10.1034/j.1600-065x.2001.1840113.x (2001).

25 Sahin, K. et al. Effect of chromium on carbohydrate and lipid metabolism in a rat model of type 2 diabetes mellitus: the fat-fed, streptozotocin-treated rat. Metabolism 56, 1233–1240, doi:10.1016/j.metabol.2007.04.021 (2007).

26 Saeedi, P. et al. Global and regional diabetes prevalence estimates for 2019 and projections for 2030 and 2045: Results from the International Diabetes Federation Diabetes Atlas, 9(th) edition. Diabetes Res Clin Pract 157, 107843, doi:10.1016/j.diabres.2019.107843 (2019).

27 Kushner, J. A. The role of aging upon beta cell turnover. The Journal of clinical investigation 123, 990–995, doi:10.1172/JCI64095 (2013).

